# Human uniqueness illustrated by life history diversity among small-scale societies and chimpanzees

**DOI:** 10.1101/2020.09.02.280602

**Authors:** Raziel J. Davison, Michael D. Gurven

## Abstract

**Background:** We compare life histories and selection forces among chimpanzees and human subsistence societies in order to identify the age-specific vital rates that best explain fitness variation, selection pressures and species divergence.

**Methods:** We employ Life Table Response Experiments that quantify vital rate contributions to population growth rate differences. Although widespread in ecology, these methods have not been applied to human populations or to look at species differences among humans and chimpanzees. We also estimate correlations between vital rate elasticities and life history traits to investigate differences in selection pressures and test predictions of life history theory.

**Results:** Chimpanzees’ earlier maturity and higher adult mortality drive species differences, whereas infant mortality and fertility variation drive differences among humans. Human fitness is decoupled from longevity by postreproductive survival, while chimpanzees forfeit higher potential lifetime fertility due to adult mortality attrition. Infant survival is often lower among humans, but lost fitness is recouped via short birth spacing and high peak fertility, thereby reducing selection on infant survival. Lastly, longevity and delayed maturity reduce selection on child survival, but among humans, recruitment selection is unexpectedly highest in longer-lived populations, which are also faster-growing due to high fertility.

**Conclusion:** Humans differ from chimpanzees more because of delayed maturity and adult mortality than child mortality or fertility rates. In both species, high child mortality reflects bet-hedging costs of quality/quantity tradeoffs borne by offspring, with high and variable child mortality likely regulating human population growth over evolutionary history. Among human subsistence societies, positive correlations between survival and natural fertility lead selection pressures in human subsistence societies to differ from modern populations undergoing demographic transition, due in part to positive correlations between longevity and natural fertility and negative correlations between recruitment elasticity and reproductive effort.

## Introduction

Humans and chimpanzees, whose recent common ancestor dates to 4-8 million years ago [1, 2], share behavioral adaptations and life history traits that distinguish them among other primates [3, 4]. Except for menopause, which appears unique among mammals apart from a few toothed whale species [5], human fertility is similar to chimpanzees [6], and mortality profiles of modern hunter-gatherers are closer to chimpanzees than they are to today’s lowest-mortality industrialized populations [7]. Nevertheless, there is great variation among human and chimpanzee life histories [8]. Here, we characterize human uniqueness by identifying the vital rates that drive life history differences within- and between-species. We interpret population life histories in terms of the slow-fast life history continuum [9] and characterize the tradeoffs that likely shaped human life history evolution [10].

Whereas many contemporary small-scale societies are growing rapidly [11], documented chimpanzee populations are typically shrinking, though favorable conditions promote increase in some wild groups [12, 13]. With the largest high-quality dataset assembled to date on fertility and mortality among human subsistence societies and chimpanzees [14], we employ life table response experiments [LTREs, 15] to quantify the importance of particular age classes for driving population growth and identify the vital rates that best explain the divergence of human and chimpanzee life histories.

Although demographic patterns are well-described for many primate species, including chimpanzees [16-18], we provide a comprehensive, up-to-date and timely comparison of human and chimpanzee life histories through the lens of the elasticities and fitness contributions of survival and fertility at different ages. These fitness contributions illustrate how life history event schedules drive differences within and between species, while elasticities reflect the force of selection and highlight the potential for fitness contributions if vital rates vary across populations [19]. Previous comparisons used fewer populations, focused primarily on composite fertility and longevity differences, and did not systematically quantify species life history differences [8, 17]. Because species comparisons are complicated by within-species life history variation [17], we compare species mean life histories and we also characterize the population drivers within each species. Using the average hunter-gatherer life history (HG) as a common reference, we compare hunter-gatherers with other natural fertility subsistence societies and with seven chimpanzee populations, including captive and managed populations representing chimpanzee “best case” life histories. We also compare the hunter-gatherer reference to three other composite life histories representing the average life histories calculated across non-exclusive foragers, and across chimpanzee populations exhibiting decreasing vs. increasing population growth. Without relying on model life tables or indirect demographic methods for age assignment or vital rate estimation, our dataset includes ten small-scale societies (five hunter-gatherers, three forager-horticulturalists, a pastoralist society and an “acculturated” hunter-gatherer population with some food production) and seven chimpanzee populations with high quality fertility and mortality data (five wild, one managed and one captive).

We identify the vital rates that are most important in driving population growth and decline using fixed LTREs [15], which decompose individual vital rate contributions to observed differences in population growth rates. We compare these results with the more widely-used elasticity analyses that prospectively estimate the potential fitness effects of vital rates [15, 19]. Differences between realized fitness contributions and the potential suggested by elasticities may indicate constraints on life history evolution.

In this context, we evaluate three predictions based on fitness elasticities: (**P1**) juvenile recruitment should be the most important for population growth [20]; (**P2**) intrinsic population growth rates (*r*) should be greater in populations with higher life expectancy (*e*_*0*_) and with higher total fertility rate (*TFR*); **(P3)** elasticity to child survival should be negatively correlated with life expectancy but positively with fertility [21].

**P1** relies on the high elasticities of infant and child survival, which are larger than elasticities to adult survival or fertility, to predict that recruitment will be most important for population fitness differences [20]. However, if stabilizing selection reduces variation in important vital rates [22], fitness contributions of high-elasticity rates are likely to be small [23]. More generally, when elasticities overestimate fitness effects this may reflect constraints on stabilizing selection that would otherwise reduce variation in these important vital rates, whereas underestimation implies that vital rate differences are more important than *a priori* predictions from the force of selection.

**P2** is the intuitive prediction that population growth should reflect both survival and reproduction, since either will increase population growth, all else equal. However, longevity and fertility may trade off [24-26]. For instance, greater life expectancy is associated with lower fertility across modern industrial nations [27], driving a negative correlation between life expectancy and population growth [28]. Therefore, the degree (and even the sign) of the correlations of population growth with fertility vs. longevity are empirical questions that we answer in the case of natural fertility subsistence populations and chimpanzees.

**P3** arises as a consequence of the slow life history of primates [29]. When infant mortality is low, more survive to maturity, thereby reducing selection on recruitment. A longer lifespan also permits replacement of dead offspring with new births, while low fertility raises the average age of a population. Because all of these effects make newborn survival less important to population fitness, elasticity to child survival is predicted to correlate negatively with life expectancy but positively with fertility (P3, [21]). We extend this logic to predict that recruitment elasticity should also correlate positively with the pace of fertility and thus negatively with mean age at first birth (*AFB*), mean age of childbearing (*MAC*) and inter-birth intervals (*IBI*) since smaller values increase fertility, but positively with age at last birth (*ALB*).

## Materials and Methods

### Demographic Data

We examine fertility and mortality rates published for ten contemporary, non-industrial small-scale societies with natural fertility and minimal to no access to modern medicine during the period corresponding to the demographic data (S1, S2 Tables; S1 File contains ethnographic details): Australian Aborigines (Northern Territory, Australia), Ache (Paraguay), Agta (Philippines), Gainj (Papua New Guinea), Hadza (Tanzania), Herero (Namibia), Hiwi (Venezuela), Ju/’hoansi !Kung (Botswana and Namibia), Tsimane (Bolivia) and Yanomamo (Venezuela and Brazil). We also examine seven chimpanzee populations, including five wild populations at Gombe and Mahale (Tanzania), Kanyawara and Ngogo (Uganda), and Taï (Ivory Coast), a captive population in the Taronga Zoo (Sydney, Australia), and a reintroduced (captive-founded but wild-breeding) population in Gambia. These captive and managed populations are not included in species-level comparative statistics or composite life histories, but are used to reflect “best-case” scenarios for chimpanzees: low mortality in the protected and provisioned Gambia population and high fertility in the captive breeding program at Taronga Zoo. Because fertility estimates for Ngogo chimpanzees are not published, we estimate contributions applying fertility estimated at nearby Kanyawara. Also, because the Taronga Zoo mortality data includes few chimpanzee deaths we use mortality data averaged across three zoo populations [30].

We employ parametric models of mortality and non-parametric models of fertility to obtain smoothed annual rates (see S1 File for details). Briefly, Siler’s [31] five-parameter competing hazard model of mortality jointly models juvenile, age-independent and adult mortality. The Siler model, estimated here with the NLIN procedure in SAS 9.4, was employed in previous treatments of human subsistence and chimpanzee mortality because of its simplicity, robustness and interpretability of its parameters [32, 33]. Using the statistical software *R* (version 3.5.1), we smooth raw fertility data with a local polynomial regression (loess; span = 0.5) and constrain the smoothed data to the observed ages of reproduction by heavily weighting zero values in the single-year age-classes before the minimum age at first birth and after the last recorded birth, and imputing values outside this range as zero. Resulting smoothed fertility was rescaled evenly across age to conserve the *TFR* from raw data (S1 Fig) and statistical predictions of *AFB* and *ALB* are close to those of source estimates (S2 Fig).

### Data Analysis

We construct a female age-structured Leslie [34] population projection matrix **A** (**A** = {*a*_*ij*_}) where matrix elements *a*_*ij*_ describe the number (*n*_*i*_) of age *i* individuals alive in the population at time *t*+1 that are contributed by one age *j* individual alive at time *t*, either via survival (*a*_*x+1,x*_ = *p*_*x*_) or fertility transitions (*a*_*1x*_ = *m*_*x*_) ([15]; S2 Table contains variable definitions; S1 File contains details of matrix model methods and calculations of life history traits). Population size is updated by applying the population projection matrix **A** to the population age structure **n** (**n** = {*n*_*i*_}) and stable asymptotic population growth is described by the dominant eigenvalue *λ* (**n**(*t*+1) = **A n**(*t*) = *λ* **n**(*t*)). From the matrix **A** we calculate vital rate sensitivities (*s*_*ij*_) reflecting the force of selection on a vital rate as well as elasticities (*e*_*ij*_) scaling the proportional effect on population growth (*e*_*ij*_ = (∂ *λ* / ∂ *a*_*ij*_) = *s*_*ij*_ (*λ* / *a*_*ij*_) [15, 19]. Because elasticities conveniently sum to unity (1 = **S**_***i***,***j***_ *e*_*ij*_), we can add elasticities across vital rates across age *x* to estimate the total elasticity to survival (*E*_*s*_ = **S**_*x*_ *e*_*x+1,x*_) or to fertility (*E*_*f*_ = **S**_*x*_ *e*_*1x*_; 1 = *E*_*s*_ + *E*_*f*_), or sum across specific ages (e.g., before or after reproductive maturity at age *α*) to distinguish the elasticity to survival through childhood (*E*_*c*_ = **S**_*x<α*_ *e*_*x+1,x*_) vs. across adulthood (*E*_*a*_ = **S**_*x≥α*_ *e*_*x+1,x*_; *E*_*s*_ = *E*_*c*_ + *E*_*a*_).

Differences in population growth rates (*λ, r* = ln *λ*) are decomposed into positive and negative contributions (*C*_*ij*_) made by vital rate differences (Δ*a*_*ij*_) to the total difference Δ*λ* using a one-way fixed-treatment life table response experiment, or LTRE (Δ*λ* = **S**_*i,j*_ *C*_*ij*_ = **S**_*i,j*_ *s*_*j*_ Δ*a*_*ij*_; Δ*a*_*ij*_ = *a*_*ij*_ ^(*m*)^ - *a*_*ij*_ ^(R)^; [15]; S1 File). Here, each population *m* (*m* = 1, 2, 3, …, *M*) is compared to a common (composite) reference (R) life history, here exhibiting the average fertility and survival rates estimated across hunter-gatherers (labeled HG and summarized in the matrix **A**^(HG)^). Species differences are highlighted by comparing this common reference to composite life histories exhibiting vital rates averaged across all wild chimpanzees (WC), and within-species differences are summarized by results for composite life histories estimated separately for exclusive hunter-gatherers (HG) vs. non-forager (NF) subsistence populations and for increasing (WC+) vs. decreasing chimpanzees (WC-). In addition to vital rate *contributions* (*C*_*ij*_) that sum to the total difference in population growth rates (Δ*λ* = **S**_*i,j*_ *C*_*ij*_), we also examine combined *effects* (*C*_*ij\**_ = |*C*_*ij*_| / **S**_*i,j*_ |*C*_*ij*_|), which are analogous to elasticities in that they sum to unity (1 = **S**_*i,j*_ *C*_*ij\**_) and which reflect the *relative* effect of fitness contributions (e.g., survival across childhood vs. adulthood) (*C*_*c*_ = **S**_*x*<*α*_ *C*_*x+1,x\**_; *C*_*a*_ = **S**_x≥*α*_ *C*_*x+1,x\**_; *C*_*s*_ = **S**_x_ *C*_*x+1,x\**_; *C*_*f*_ = **S**_x_ *C*_*1x\**_; 1 = *C*_*s*_ + *C*_*f*_). For comparison, fertility is binned into early, prime and late fertility effects at the ages when completed fertility is 0-25%, 25-75% and 75-100% of *TFR* in the hunter-gather reference (ages 0 to 22, 23 to 35, and 36 to 50, respectively). To aid interpretation, we calculate standard demographic rates: mortality hazard (*μ*_*x*_), survivorship (*l*_*x*_), life expectancy at birth (*e*_*0*_), total fertility rate (*TFR*), mean age at first birth (*AFB*), mean age of childbearing (*MAC*), mean age at last birth (*ALB*) and mean inter-birth intervals (*IBI*) (S2 Table; S1 File contains calculations).

We also evaluate three predictions of population biology and life history theory (**P1-P3**).

**P1:** Based on their large elasticities, we expect child survival contributions to be the largest [20], but because strong stabilizing selection may canalize important rates [22], thereby limiting LTRE contributions [23], we expect child survival effects to be smaller than elasticities predict. We calculate a scalar ratio (*Z*_*ij*_) that reflects the actual realized contributions of vital rates, relative to the potential suggested by elasticities (*Z*_*ij*_ = *C*_*ij\**_ / *e*_*ij*_). Because both vital rate effects (*C*_*ij\**_) and elasticities (*e*_*ij*_) sum to unity, we can estimate *Z* across all of childhood (*Z*_*c*_ = *C*_*c*_ / *E*_*c*_) or across adulthood (*Z*_*a*_ = *C*_*a*_ / *E*_*a*_) and for lifetime fertility (*Z*_*f*_ = *C*_*f*_ / *E*_*f*_).

**P2**: Both survival and fertility should increase population growth, which would drive positive correlations between population growth rates (*r* = ln *λ*) and two emergent life history traits: life expectancy (*e*_*0*_) and lifetime fertility (*TFR*).

**P3**: Neonate mortality, which is under the strongest selection pressure (*E*_*0*_ = *e*_*21*_ = max_*ij*_ (*e*_*ij*_)), should be negatively correlated with longevity because longer lifespans increase the average age of the population, but positively correlated with fertility because higher fertility increases the proportion of the population in the youngest age-classes [21]. Since neonate survival commands a lower premium under slower life histories exhibiting longevity and low or delayed fertility, *E*_*0*_ (which is relatively high among fast life histories) should also correlate positively with the tempo of reproduction and thus negatively with *AFB, MAC* and *ALB*, but positively with *IBI*.

We report *p*-values from non-parametric Mann-Whitney-Wilcoxon rank-sum tests used for all statistical tests of differences in means; for associations we report correlation coefficients *r* and significance *p*-values. All results were computed using *Matlab* (*Mathworks, 2018b*).

## Results

### Vital Rates and Elasticities

#### Mortality

Among hunter-gatherers, predicted neonate (age 0-1) mortality is higher than among chimpanzees (increasing or declining) and infant mortality (age 1-2) is higher than among increasing chimpanzee populations, but average hunter-gatherer mortality is lower at all other ages (Fig 1A; S3 Fig). Non-foragers have lower mean mortality than hunter-gatherers except between ages 53-64, where they are equivalent. Human life expectancies in our sample are more than twice those of wild chimpanzees (*e*_*0*_; *p =* 0.005, Wilcoxon rank sum test) and are marginally higher among non-foragers than among hunter-gatherers (*p* = 0.056; Table 1, S3 Table). Captive and reintroduced populations are at the upper end of the wild chimpanzee range of longevity and several hunter-gatherer populations are at the lower end of the human range.

**Table 1.**
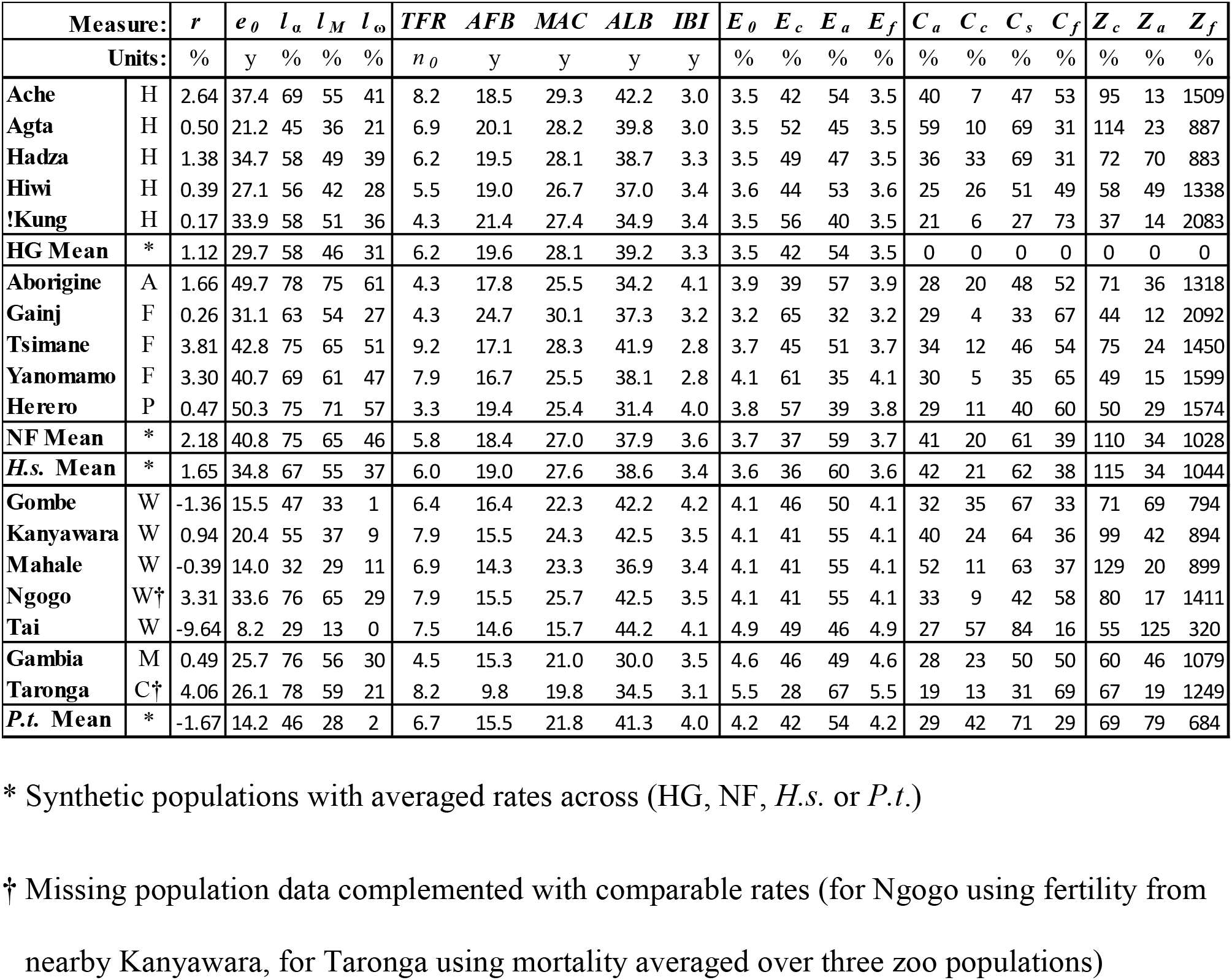
Summary of demographic measures for chimpanzee and human populations. For ten small-scale human societies and seven chimpanzee populations (appearing in rows), columns note subsistence ecology, annual growth rate (%), life expectancy (y), fertility measures (*TFR* in units of neonates *n*_*0*_), age schedules of life history events (y), proportions (%) of elasticities (*E*) and contributions (*C*) due to different life cycle components, and ratios of realized contributions to prospective elasticities (*Z* = *C*/*E*). Separate rows show results for the mean life histories of hunter-gatherers (HG Mean, the LTRE reference), non-foragers (NF Mean), human small-scale societies (H.s. Mean), and wild chimpanzees (P.t. Mean). Human subsistence ecologies in the second column are abbreviated: H (hunter-gatherer), A (acculturated hunter-gatherer), F (forager-horticulturalist) or P (pastoralist); chimpanzee management status is abbreviated: W (wild), M (managed) or C (captive).

**Fig 1.**
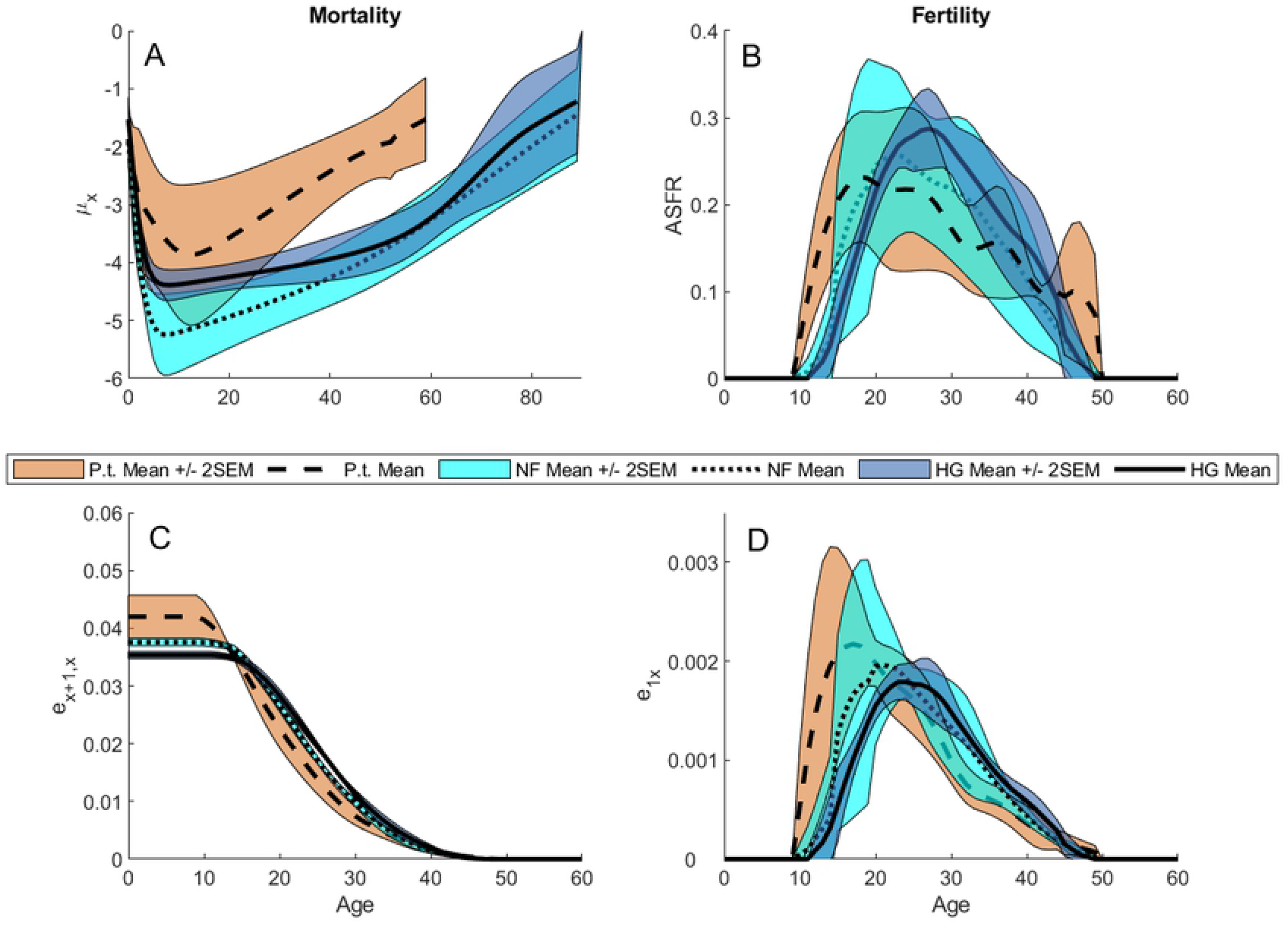
Summary statistics for vital rates and elasticities. 95% Confidence Intervals (Mean ± 2 SEM) are calculated across five hunter-gatherer societies (dark blue fill, solid lines), across five non-exclusive forager societies (light blue fill, dotted lines) or across five wild chimpanzee populations (red fill, dashed lines). (A), Mortality (*μ*_*x*_). (B), Age-specific fertility rate (*ASFR*). (C), Survival elasticities (*e*_*x+1,x*_). (D), Fertility elasticities (*e*_*1x*_).

After age 4, human mortality rates are lower than those in any chimpanzee population, but infant mortality (age 0 to 1) is higher than the chimpanzee mean in five small-scale societies (the Agta, Hadza, Hiwi, Ju/’hoansi !Kung and Yanomamo). The lowest infant mortality (age 1 to 2) in our sample is among the managed Gambia chimpanzees, and the lowest mortality between ages 2 and 4 is among wild Ngogo chimpanzees (S3 Fig). Humans are marginally more likely than chimpanzees to survive to their later age of reproductive maturity (*l*_*α*_; *p* = 0.099), but are significantly more likely to survive to the mean age of childbirth (*l*_*M*_; *p* = 0.040) and to the maximum age of reproduction (*l*_*ω*_; *p* = 0.005; Table 1, S3 Table). These species differences are driven more by non-foragers (*p* = 0.095 [*l*_*α*_]; *p* = 0.032 [*l*_*M*_], *p* = 0.016 [*l*_*ω*_]), whereas among hunter-gatherers, survivorship is lower than among non-foragers (*p* = 0.032 [*l*_*α*_]; *p* = 0.016 [*l*_*M*_]; *p* = 0.095 [*l*_*ω*_]) and only survivorship to the maximum *ALB* is significantly higher than chimpanzees (*l*_*ω*_; *p* = 0.032; S3 Table).

#### Fertility

Although mean survival-conditioned fertility (*TFR*) is similar among humans and chimpanzees (*P* > 0.1; Fig 1B, S3 Table), maximum lifetime fertility is highest among humans (Tsimane *TFR* = 9.2; Table 1) and as noted above (*l*_*ω*_ comparison), very few chimpanzees survive to complete their potential *TFR* (10% of chimpanzees, compared to 33% of hunter-gatherers and 49% of non-foragers). As noted in previous studies [6], chimpanzees have earlier mean *AFB* (*p* = 0.001) and *MAC* (*p* = 0.005) than humans, but later *ALB* (*p* = 0.037) and longer *IBI*s (*p* = 0.025) (Tables 1, S3 Table). While both hunter-gatherers and non-foragers have later *AFB* than chimpanzees (*p* = 0.008 for each), non-foragers and chimpanzees have similar *MAC* (*p* = 0.056) and *IBI* (*p* > 0.1), and hunter-gatherers and chimpanzees have similar *ALB* (*p* > 0.1). Chimpanzee interbirth intervals calculated using these *AFB* and *ALB* estimates (mean±SD *IBI* = 3.7±0.4y) exceed human *IBI*s (*p* = 0.025), but this difference is only significant for hunter-gathereres (*p* = 0.008). These chimpanzee *IBI*s are also shorter than the 5.1-6.2y intervals reported elsewhere [6, 35]. As might be expected, our *IBI* estimate falls between those calculated for mothers whose offspring died before vs. after age four (2.2y and 5.7y, respectively; [6]), with our lower estimate reflecting the averaged effects of infant mortality on birth spacing. Closer examination shows population differences in the tempo of fertility (Table 1, S3 Table).

#### Elasticities

Compared to chimpanzees, human elasticity to infant survival is lower (*E*_*0*_; *p* = 0.001; *p* = 0.008 [hunter-gatherers]; *p* = 0.016 [non-foragers]), but elasticities to child survival (*E*_*c*_; *p* = 0.099; *p* = 0.095 [hunter-gatherers]; *p* > 0.1 [non-foragers]) and to adult survival are similar to chimpanzees (*E*_*a*_; *p* > 0.1), and total elasticity to human fertility is lower (*E*_*f*_; *p* = 0.001 [humans]; *p* = 0.008 [hunter-gatherers]; *p* = 0.016 [non-foragers]) (Fig 1D, S4 Fig; Table 1, S2 Table). Fertility elasticities may climb rapidly with age (e.g., Herero, Yanomamo and Taï chimpanzees) or slowly (e.g., Ache, Hadza and Tsimane) depending on the pace of fertility, but decrease at approximately the same rate as survival elasticities due to mortality attrition affecting both simultaneously (Fig 1C,D).

### Fitness Contributions

All ten small-scale societies and two wild chimpanzee populations are growing, but two chimpanzee groups are declining slowly and one is collapsing (Fig 2; Table 1). However, due to wide variation among our small sample, population growth differences are not statistically significant (*r* = log *λ, p* > 0.1; S1 Table). Compared to the hunter-gatherer reference, lower neonate (age 0) mortality elevates population growth among both increasing and declining chimpanzees, but lower mortality at other ages makes larger negative contributions.

**Fig 2.**
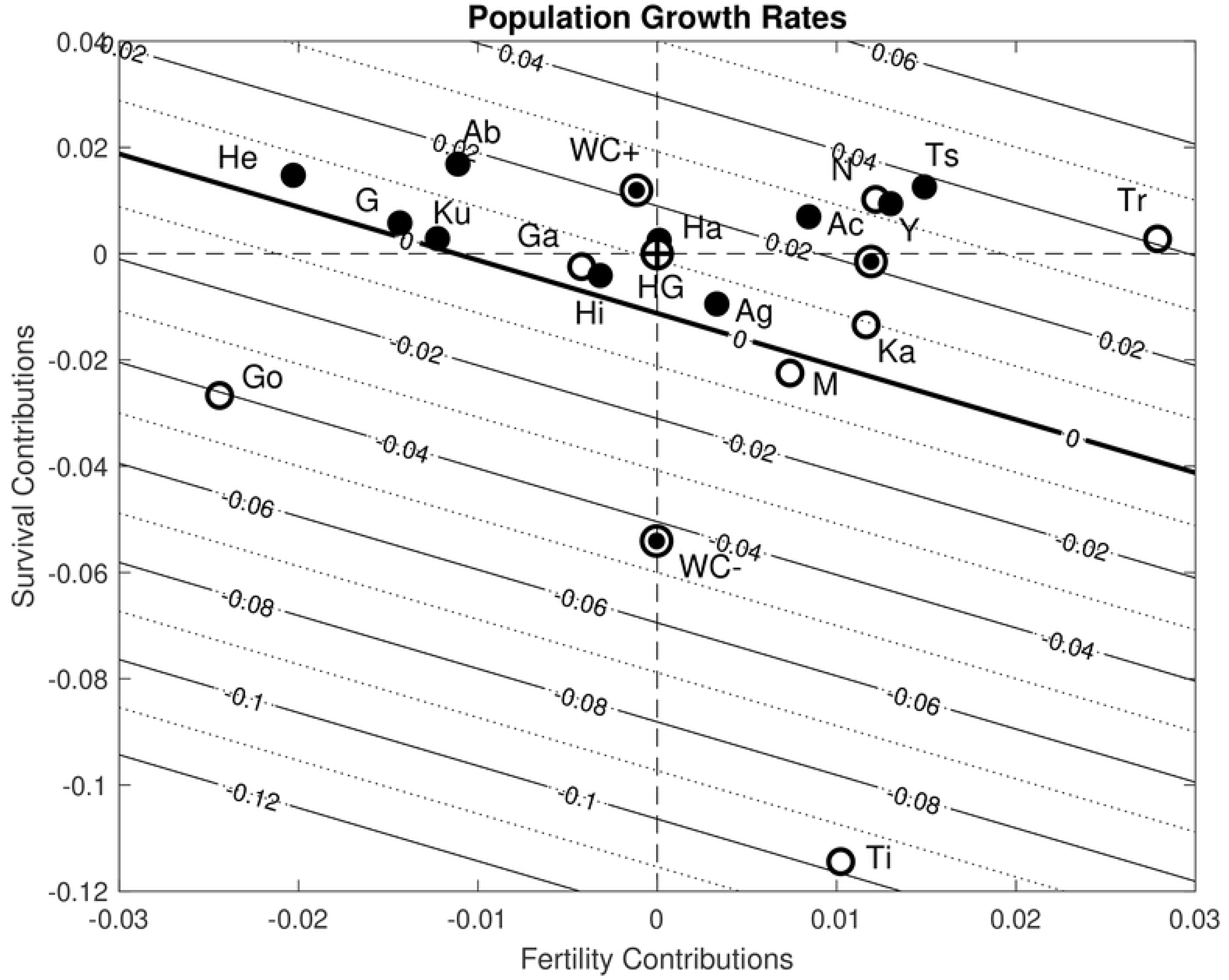
Net contributions of fertility (*x*-axis) and survival (*y*-axis) from a Life Table Response Experiment (LTRE) comparing humans and chimpanzees to mean life history estimated across five hunter-gatherer societies. Hunter-gatherer societies are indicated by filled circles, non-foragers by filled squares, and chimpanzees by open circles; the non-forager mean life history (labeled NF) and the mean life histories for declining (WC-) and increasing (WC+) chimpanzees are each indicated with a black-and-white dot, and the mean hunter-gatherer (HG) reference by a large black bullseye at the origin. Compared to the HG reference, populations have higher survival if they fall above the horizontal dashed line and higher fertility if they fall to the right of the vertical dashed line. Humans are labeled: Ab (Aborigines), Ac (Ache), Ag (Agta), G (Gainj), Ha (Hadza), He (Herero), Hi (Hiwi), Ku (Ju/’hoansi !Kung), Ts (Tsimane), Y(Yanomamo); chimpanzee populations are labeled: Ga (Gambia), Go (Gombe), Ka (Kanyawara), N (Ngogo), Ti (Taï), Tr (Taronga).

Positive contributions of chimpanzees’ higher early fertility up to age 22 (age 29 at Kanyawara) are partially offset by lower prime-age fertility between ages 23 and 35, which comprises half of the mean human *TFR* (Figs 2B, 3), but higher survival allows positive net contributions of late fertility among increasing but not decreasing chimpanzees. The rapid population growth of non-foragers is mainly due to higher survival at all ages, but offset by prime and late-age fertility, which are lower than the hunter-gatherer reference (Fig 3).

**Fig 3.**
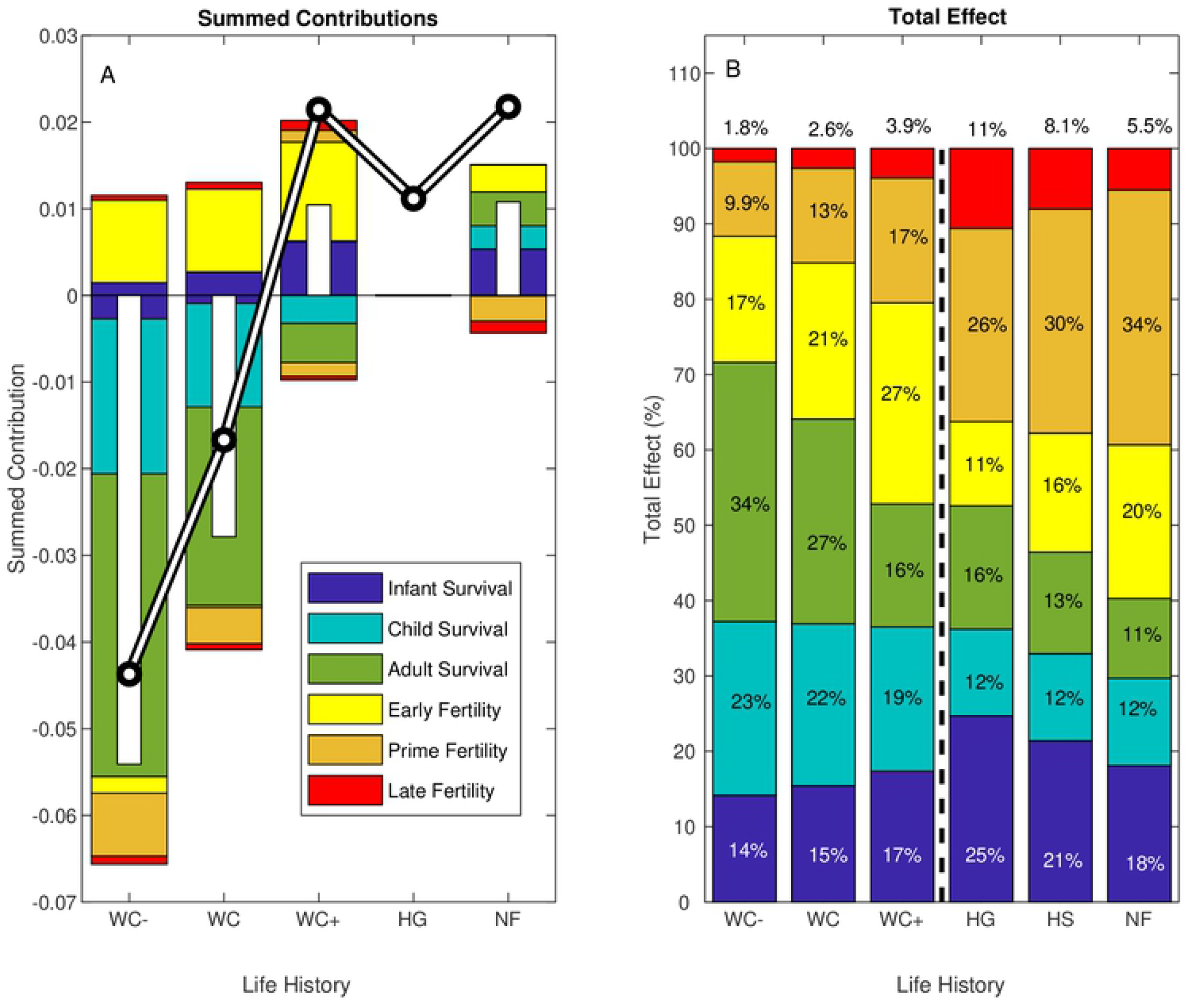
Summed contributions for each composite life history. (A), stacked bars show summed contributions of infant, child and adult survival and of early, prime and late fertility. Positive and negative contributions are summed separately to reflect the net difference in population growth rate (white bars), relative to the composite mean hunter-gatherer reference (note zero contributions for HG); the black-and-white line crossing the bars indicates the population growth rate (*r* = log(*λ*)). Results are shown for four composite life histories with vital rates averaged over: declining chimpanzees (WC-), all wild chimpanzees (WC), increasing chimpanzees (WC+), hunter-gatherers (HG), or non-foragers (NF). (B), total effects (**S** *C*_*ij*\*_) reflecting the combined magnitude of contributions, are averaged across the populations within each of the groupings in *A* plus the average across all human (HS). Stacked bars decompose the mean total effect (the proportion of the combined magnitude of all contributions) made by infant, child and adult survival and by early, prime and late fertility (inset text shows the percent of total effects, with late fertility effects labeled above the bars).

Despite differences in the signs of fertility and survival contributions, the relative magnitudes of vital rate effects are similar across composite life histories. The only significant difference between hunter-gatherers, non-foragers and chimpanzees is that survival effects are stronger among chimpanzees than non-foragers (*C*_*s*_; *p* = 0.032) and fertility effects are stronger among non-foragers than among hunter-gatherers (*C*_*f*_; *p* = 0.032; Fig 3B, S3 Table).

Because so few chimpanzees survive to advanced ages, large differences in the potential for late-life fertility contribute little to population growth. Among humans, high survival drives population growth among non-foragers; among hunter-gatherers, lower early fertility effects are offset by higher prime- and late-age fertility (Fig 3A; S5, S7 Figs).

Among chimpanzees, slow decline at Gombe was close to the rate calculated for the mean chimpanzee life history, whereas decline at Mahale was slower despite high infant mortality because of higher survival and early fertility. Population growth at Kanyawara and Ngogo were due to lower mortality and higher prime fertility (S6, S7 Figs). At Taï, high child and adult mortality drove precipitous decline (*r* = -9.6%) despite low infant mortality and fertility near the chimpanzee mean. The managed population at Gambia was near-stationary with longevity balancing low fertility and the Taronga Zoo population was growing rapidly with an active breeding program.

Although human fertility was mostly lower than chimpanzees, the populations with the highest growth rates (i.e. Tsimane, Yanomamo, and chimpanzees at Ngogo and Taronga Zoo) also had high fertility (S5, S7 Figs). The Ju/’hoansi !Kung, Gainj and Hiwi were all near stationary population growth – the Hiwi because of low infant survival and low fertility, whereas the Gainj and Ju/’hoansi had higher survival balanced by lower fertility (Fig 2C; S5, S7 Figs). Among the Herero, high survival at all ages offset very low fertility. The Agta and Hadza both had low early fertility, but high infant mortality drove slower growth among the Agta despite higher prime and late fertility. Relatively rapid growth among the Northern Territory Aborigines was due to high survival offsetting low fertility at all ages, whereas the Ache grew faster due to high prime and late fertility. Very rapid growth was due to survival and early fertility among the Yanomamo and due to survival and late fertility among the Tsimane.

**P1:** Survival (especially recruitment) should be the most important for fitness (*E*_*0*_ > *E*_*c*_ > *E*_*a*_ > *E*_*f*_; *C*_*0*_ > *C*_*c*_ > *C*_*a*_ > *C*_*f*_). In agreement with P1, infant mortality has the largest elasticity and in many human populations high infant mortality substantially reduced population growth relative to the chimpanzee reference. Neonate survival did make the largest contribution (*C\** = *p*_*0*_) in four out of five hunter-gatherer societies, four out of five non-foragers and three out of five wild chimpanzee populations, with infant survival (*p*_*1*_) making the largest contribution among the Hiwi and the Yanomamo and among chimpanzees at Mahale and Ngogo (Table 1). Also consistent with P1, the combined effects of neonate, infant and child survival across the life cycle were larger than adult survival effects (*C*_*c*_ > *C*_*a*_; *p* < 0.001 [humans]; *p* = 0.008 [hunter-gatherers]; *p* = 0.048 [non-foragers]; *p* = 0.008 [chimpanzees]) and larger than fertility effects in chimpanzees (*C*_*c*_ > *C*_*f*_; *p* = 0.008 [chimpanzees]). However, fertility effects were unexpectedly larger than child survival effects in humans (*C*_*c*_ < *C*_*f*_; *p* < 0.001 [humans]; *p* = 0.008 [hunter-gatherers]; *p* = 0.008 [non-foragers]; S4 Table, S8 Fig). Across all populations pooled and across hunter-gatherers alone, fertility and survival effects were equivalent (*C*_*s*_ ≈. *C*_*f*_; *p* > 0.1), but survival effects were larger among chimpanzees (*C*_*s*_ > *C*_*f*_; *p* = 0.016) and fertility effects were larger among non-foragers (*C*_*s*_ < *C*_*f*_; *p* = 0.008; S4 Table).

Elasticities estimate the force of selection and reflect the *potential* for fitness effects if vital rates differ. However, the actual (realized) effects of vital rate differences depend on *observed* population-level differences, and the effect:potential ratio may inform us about the tradeoffs constraining life history evolution. Although elasticities similarly overestimate the fitness importance of child vs. adult survival among hunter-gatherers (*Z*_*c*_ ≈ *Z*_*a*_; *p* > 0.1), child survival effects were underestimated more among chimpanzees (*Z*_*c*_ < *Z*_*a*_; *p* = 0.016) and (marginally) among non-foragers (*Z*_*c*_ < *Z*_*a*_; *p* = 0.095). Fertility effects were grossly underestimated, within and between species (*Z*_*c*_ << *Z*_*f*_; *p* = 0.008; S4 Table).

**P2:** Population growth rates should be higher with longevity and high fertility. Consistent with P2, population growth (*r*) is positively correlated with life expectancy (*e*_*0*_) across our two-species sample (corr(*r, e*_*0*_); *r* = 0.67, *p* = 0.006) and (marginally) across chimpanzee populations (*r* = 0.83, *p* = 0.084), while across human populations population growth and fertility are positively correlated (corr(*r, TFR*); *r* = 0.81, *p* = 0.004; *r* = 0.95, *p* = 0.014 [non-foragers]; *r* = 0.82, *p* = 0.090 [hunter-gatherers]; Table 2). Inconsistent with P2, population growth is not correlated with life expectancy across humans or with fertility across chimpanzees (*p* > 0.1).

**Table 2.** Life history correlations. Rows show correlations of: (P2) population growth rates (*r*) with life expectancy (*e*_*0*_) or fertility (*TFR*); (P3) elasticity to child survival (*E*_*0*_) with life history traits (*e*_*0*_, *TFR*). Columns indicate statistics (*r* coefficients and *p*-values) for correlations across all populations pooled, across wild chimpanzee populations, across small-scale human societies, across hunter-gatherers, or across non-foragers. Greyed cells indicate NS correlations and lightly greyed cells indicate marginal significance (0.05 < *p* < 0.1). Bold values indicate deviations from predictions (P3).

|  |  | <u>All</u> |  | <u>Chimpanzees</u> |  | <u>Humans</u> |  | <u>Hunter-Gatherers</u> |  | <u>Non-Foragers</u> |  |
| --- | --- | --- | --- | --- | --- | --- | --- | --- | --- | --- | --- |
| | | $r$ | $p$ | $r$ | $p$ | $r$ | $p$ | $r$ | $p$ | $r$ | $p$ |
| P2 | $r \ e_0$ | 0.671 | 0.006 | 0.827 | 0.084 | 0.417 | 0.231 | 0.623 | 0.262 | 0.078 | 0.900 |
| | $r \ TFR$ | 0.125 | 0.659 | 0.200 | 0.747 | 0.811 | 0.004 | 0.820 | 0.090 | 0.948 | 0.014 |
| P3 | $E_0 \ e_0$ | -0.411 | 0.051 | -0.572 | 0.313 | **0.686** | **0.029** | 0.112 | 0.858 | 0.734 | 0.158 |
| | $E_0 \ TFR$ | 0.296 | 0.170 | 0.078 | 0.901 | 0.116 | 0.751 | -0.396 | 0.509 | 0.278 | 0.651 |
| | $E_0 \ AFB$ | -0.897 | $< 10^{-3}$ | -0.392 | 0.515 | -0.827 | 0.003 | -0.296 | 0.629 | -0.927 | 0.023 |
| | $E_0 \ MAC$ | -0.885 | $< 10^{-3}$ | -0.948 | 0.014 | -0.875 | 0.001 | -0.683 | 0.203 | -0.910 | 0.032 |
| | $E_0 \ ALB$ | -0.058 | 0.792 | 0.549 | 0.338 | -0.295 | 0.409 | -0.408 | 0.496 | -0.161 | 0.796 |
| | $E_0 \ IBI$ | 0.293 | 0.174 | 0.607 | 0.278 | 0.254 | 0.479 | **0.814** | **0.094** | 0.148 | 0.813 |

**P3:** Longevity should decrease, and fertility increase, the fitness elasticity to recruitment (corr(*E*_*0*_, *e*_*0*_) < 0; corr(*E*_*0*_, *TFR*) > 0). Consistent with P3 [21], elasticity to child survival is always the largest elasticity and is negatively correlated (marginally) with life expectancy across species (corr(*E*_*0*_, *e*_*0*_); *r* = -0.41, *p* = 0.051; Table 2), but not across chimpanzees (*p* > 0.1). Inconsistent with P3, *E*_*0*_ is *positively* correlated with *e*_*0*_ across humans (*r* = 0.69, *p* = 0.029), and there is no correlation between *E*_*0*_ and *TFR* within or across species (*p* > 0.1; Table 2). Across our pooled two-species sample we do find predicted (P3) negative correlations of *E*_*0*_ with *AFB* (*r* = -0.90, *p* < 0.001) and *MAC* (*r* = -0.89, *p* < 0.001), but not with *ALB* (*p* > 0.1). Among chimpanzees alone, only *MAC* is negatively correlated with *E*_*0*_ (*r* = -0.95, *p* = 0.014); among humans, *E*_*0*_ is negatively correlated with *AFB* (*r* = -0.83, *p* = 0.003; *r* = -0.93, *p* = 0.023 [non-foragers]; *p* > 0.1 [hunter-gatherers]) and *MAC* (*r* = -0.88, *p* = 0.001; *r* = -0.91, *p* = 0.032 [non-foragers]; *p* > 0.1 [hunter-gatherers]; Table 2) but not with *ALB* or *IBI* (*p* > 0.1). Among hunter-gatherers there is a (marginal) *positive* correlation between *E*_*0*_ and *IBI* (*r* = 0.81 *p* = 0.094).

## Discussion

Although elasticities usually identify survival, especially juvenile survival, as the most important vital rate for fitness (P1, [20]) other rates may have large effects. Juvenile survival was an important driver of population-and species-level differences (33% of all effects across human populations and 37% across chimpanzees), but adult survival was also an important driver (14% of all effects across humans and 27% among chimpanzees; Table 1). However, fertility contributions were two orders of magnitude greater than expected based on the elasticities reflecting their potential, and fertility played a large role in regulating the five populations nearest stationarity (four out of five hunter-gatherer groups and one foraging-horticulturalist group): low fertility balanced longevity in five populations and high late-life fertility compensated for infant mortality among the Agta. Also, high fertility drove rapid increase in the fastest-growing populations (Yanomamo, Ngogo, Tsimane and Taronga). That we found such large contributions of fertility differences (54% of all effects among humans and 36% among chimpanzees; Table 1) highlights the potential for low-elasticity vital rates to have large effects on population fitness if they differ more than high-elasticity rates [36]. This counterintuitive result is what we should expect if stabilizing selection canalizes important vital rates [22, 23]. The higher early fertility of non-foragers nearly balanced the higher prime and late fertility of hunter-gatherers, and among non-foragers these opposing fertility effects were larger than survival effects. This highlights a valuable feature of LTRE contributions, which allow us to identify the vital rates driving opposing fitness effects even when their signs and magnitudes balance to drive small net contributions.

We report several findings suggesting constraints on the evolution of slower human life histories. Child survival among small-scale societies overlaps with rates documented for chimpanzees and child survival varies much more across populations than adult survival. Lower variation across populations in adult human mortality estimates may reflect greater buffering of extrinsic mortality through derived human traits like food storage, widespread food sharing and ethnomedicine. Higher variation in chimpanzee mortality may reflect transient dynamics causing chimpanzee declines over the past century, due in part to human impacts such as poaching, habitat destruction and infectious outbreaks [37]. Despite strong stabilizing selection, child survival also varies over time more than adult survival among humans [38] and among non-human primates [39]), reflecting greater juvenile vulnerability to environmental effects. Because of quality-quantity trade-offs in which high fertility often comes at the expense of infant survival [40, 41], low and variable infant survival may reflect costs of reproduction borne more by offspring than adults [25, 42], and these tradeoffs may limit demographic buffering through variance-reduction in this important vital rate. Higher infant mortality among some human societies may also reflect the costs of short inter-birth intervals and overlapping child dependence. These conspicuous features of human life histories combine elements of slow life histories (late maturity and low adult mortality) with elements of a faster life history (high infant mortality and short inter-birth intervals), which are made possible through adult production surpluses and intergenerational transfers [8, 43]. Recruitment may suffer during resource shortages but indirect contributions of production transfers buffer child mortality effects [44, 45]. Also, if negative fitness effects of stochastic environments exceed costs of reproduction, then high fertility may bet-hedge against child mortality [46], resulting in higher long-term recruitment than a conservative slow life history strategy that buffers child mortality by reducing fertility [47].

As predicted (P2), population growth rates were positively correlated with longevity across our two-species sample, but they were not correlated with fertility. Within species, population growth is decoupled from longevity among small-scale subsistence societies and from fertility among chimpanzees. Among humans this reflects a slow life history and long post-reproductive lifespan, during which direct fitness contributions are zero even if individuals contribute to the fitness of living offspring indirectly through grandparenting [44] and other types of intergenerational resource transfers [45]. In contrast, high chimpanzee adult mortality decouples fitness from potential fertility because the potential contributions of higher late-life fertility are largely forfeited due to mortality attrition (only 10% of chimpanzees survive to attain their potential *TFR*, compared to 33% across hunter-gatherers and 49% across non-foragers).

Among chimpanzees, fertility contributions reflect recent high estimates of wild chimpanzee fertility (mean *TFR* = 6.7) based largely on a published compilation [6]. This survival-conditioned *TFR* is much greater than earlier estimates of 3.4 based on fewer populations and fewer births [8, 44, 48]. Those earlier estimates under-estimated the mean age at last birth (*ALB* = 27.7 y vs. our estimate of 41.3 y) and may have also under-estimated mean age of first birth due to differences between dispersing and non-dispersing females [49]. To our knowledge, the finding that potential fertility in chimpanzees is comparable with subsistence humans has not been widely appreciated, including the paper from which the fertility data originate [6]. If adult mortality rather than fertility limits chimpanzees’ reproductive potential, then human and chimpanzee life histories would be even more similar under conditions of low adult mortality. *IBI*s in chimpanzees are lower when infants die early, and so average *IBI* is affected by early life survival [14, 35]. Unlike humans, lower juvenile mortality would only lengthen chimpanzee *IBI*s, and therefore reduce lifetime fertility. Our finding that the long-lived populations have shorter *IBI*s and higher *TFR* suggests that *IBI* differences among chimpanzees are more due to ecological conditions favoring both fertility and survival rather than tradeoffs between fertility and infant survival. Low mortality at Kanyawara, and especially at Ngogo, demonstrates the potential for rapid chimpanzee population growth, with vital rates that are not too dissimilar from those of some hunter-gatherers. On the other hand, the captive zoo population illustrates the most favorable conditions, where chimpanzees have the reproductive potential to increase as rapidly as the fastest-growing human subsistence populations. While phylogenetic analysis shows that human uniqueness stems from longevity and short birth spacing more than age at maturity [51], we find that age at maturity interacts with adult mortality to drive species life histories apart by limiting prime-age fertility contributions among chimpanzees.

As predicted (P3) by Jones [21], longevity marginally eases selection on recruitment across our two-species sample, but child survival is at a greater premium among long-lived human populations, which in our sample also exhibit high fertility and rapid population growth. It is likely that the correlations Jones [21] predicted were due to cultural practices driving greater negative co-variation between fertility and mortality in Coale-Demeney model life tables than among subsistence societies (his small sample included examples with modern contraception driving low fertility and modern medicine driving low mortality). Although recruitment selection was not correlated with fertility in our sample, it was negatively correlated with fertility up to the mean age of childbearing, suggesting that the onset and peak of fertility moderate the fitness importance of recruitment more than fertility completion or birth spacing. Also, the positive correlation between hunter-gatherer recruitment elasticity and inter-birth intervals suggests that recruitment is more important when reproductive effort is low, with longer *IBI*s putting a premium on infant survival and short *IBI*s allowing replacement of lost offspring.

### Study Limitations

Our sample is the largest to date for human subsistence populations and wild chimpanzees, but these populations in their recent environments may not accurately represent ancestral life histories. The circumstances surrounding subsistence lifestyles and ecology vary by geography, extent of interactions with neighboring populations, governmental intervention and regulation of territory, and other factors. Although contemporary hunter-gatherers do not replicate ancestral demography even in these earliest recorded periods, they are the best reflection of the evolutionary context within which our species evolved and they exhibit characteristics common across prehistory among small-scale societies, including natural fertility, non-market livelihoods, greater pathogen burden, and multi-generational cooperation.

Though imperfect representatives of the past, the differences we observe within and between species nonetheless offer a unique opportunity to learn about the forces shaping human life history. Previous findings suggest that alternative demographic routes to human persistence are reflected in life history adaptations that maintain the potential for high intrinsic growth rates and allow recovery from periodic population crashes [14]. The Ju/’hoansi !Kung, Hiwi and Gainj, with low and delayed fertility, hover near-stationarity and are on the slower side of a life history continuum, while the Tsimane and Yanomamo are on the faster side with early and high fertility driving rapid population growth, perhaps in response to post-colonization recovery. The Hadza life history is close to the hunter-gatherer composite reference, suggesting perhaps that they may best represent the “typical” contemporary hunter-gatherer population. While the !Kung belong to the most ancient (L1) human haplogroup [51], their lower population growth may reflect habitat degradation and displacement by pastoralists [52]. Similarly, data on extant chimpanzees reflect novel anthropogenic influences but provide the best representation of the demography of ancestral hominins [53], with captive and managed populations providing additional insights about best-case scenarios. As with any pair of lineages, we are confronted with questions about the conditions under which human and chimpanzee life histories diverged, since they may have faced very different selection pressures over evolutionary time since their divergence from a common ancestor. Also, because we are sampling human societies that survived contact, our sample may over-represent growing populations especially since these short-term data may have captured transient growth periods in population cycles with rapid declines and prolonged recovery [14].

## Conclusions

Since divergence from chimpanzee-like ancestors, human survival has increased so much that even pre-industrial human and chimpanzee mo**r**tality profiles barely overlap. While species differences in adult mortality have been widely recognized [17], we report additional species differences and similarities: hunter-gatherers have similar, and sometimes higher, infant mortality than chimpanzees, while fertility is much more variable across human societies and overlaps the range of chimpanzees, especially across prime childbearing years. However, due to high mortality attrition, the force of selection on chimpanzee fertility is much lower than for humans and more strongly favors younger mothers.

Our findings suggest that the trajectory forward from the life history of our most recent common ancestor with the chimpanzee was likely not a monotonic decline in mortality and that high and variable infant mortality likely played a large role in regulating population growth over evolutionary time. We also find that fertility differences have substantial effects on population growth despite low elasticities, and that older individuals may contribute more to population-level fitness differences than younger individuals with higher reproductive values. The diverse environments humans inhabit are partly responsible for observed variation in reproductive success across populations, but quality-quantity tradeoffs between fertility and juvenile survival, combined with prolonged juvenile susceptibility, may constrain evolution of slower human life histories in subsistence societies with natural fertility. Because delayed fertility reduces selection on recruitment across species and among humans, this indicates a fast-slow continuum of life history even among extant hominins, with early *AFB* and strong recruitment selection on the fast side and late *AFB* and weaker recruitment selection on the slow side. High and variable juvenile mortality also reflects bet-hedging costs of reproduction, maintaining a high selective premium on juvenile survival even in longer-lived human populations. We find that late-life fertility is an important driver of population-level differences among small-scale societies despite typically low survival to these ages, and that longevity can maintain stationary populations despite low fertility. Age-patterns of mortality strongly mediate the effects of fertility differences, with adult mortality, age at maturity and menopause driving human and chimpanzee life histories apart despite similar survival-conditioned fertility.

## Supporting information

**S1 Fig. Fertility smoothing comparison (raw vs. smoothed)**. Fertility smoothed with a local polynomial regression (loess; span 0.5, blue lines) is compared with raw *ASFR* from source literature (see table 1). Note that for the Hadza, published rates are already smooth.

**S2 Fig. Fertility tempo comparisions**. (A), Estimated mean ages at first birth (*AFB*) are compared with literature values. (B), Esimated age at last birth (*ALB*). Our estimates using fertility (*ASFR*) and survivorship (*l*_*x*_) are on the *x*-axes and literature values (sources listed in table 1) are on the *y*-axes. Dashed lines show 1:1 parity line, solid lines show significant linear regressions and inset text reports correlation coefficients (*r*^*2*^) and significance *p*-values.

**S3 Fig. Mortality and fertility rates for individual populations**. Mortality *μ*_*x*_ (blue lines, left *y*-axes) and fertility *ASFR* (red lines, right *y*-axes) are shown for each age *x* (*x*-axes). Black dots indicate neonate mortality (*μ*_*0*_). Vital rates are estimated for ten small-scale human subsistence societies and seven chimpanzee populations (parenthetical labels indicate ecology type as in table 1). Bottom panels (T-W) show results for the mean life histories calculated across hunter gatherers (T), non-foragers (U), wild chimpanzees (V), declining chimpanzees (O), increasing chimpanzees (S), or all human small-scale societies (W).

**S4 Fig. Vital rate elasticities**. Elasticity to survival (*E*_*s*_, blue lines, left *y*-axes) and to fertility (*E*_*f*_, red lines, right *y*-axes) are shown for each age *x* (*x*-axes). Elasticities are estimated for ten small-scale human subsistence societies and seven chimpanzee populations (parenthetical labels indicate ecology type as in table 1). Bottom panels (T-W) show results for the mean life histories calculated across hunter gatherers (T), non-foragers (U), wild chimpanzees (V), declining chimpanzees (O), increasing chimpanzees (S), or all human small-scale societies (W).

**S5 Fig. LTRE contributions among small-scale societies**. Using average vital rates of hunter-gatherers as a reference, differences in population growth rate (*λ*) are decomposed into contributions from survival (*C*_*s*_) and fertility (*C*_*f*_) for ten small-scale human socieities, arranged by subsistence type (as in Table 3). (C), inset text reports negative neonate (age 0) survival (*p*_*0*_) contribution exceeding the *y*-axis limits.

**S6 Fig. LTRE contributions among chimpanzees**. Using average vital rates of hunter-gatherers as a reference, differences in population growth rate (*λ*) are decomposed into contributions from survival (*C*_*s*_) and fertility (*C*_*f*_) for seven populations of chimpanzees, including five wild populations (labeled W), one managed population founded by released captives (labeled M) and one captive population (labeled C). (D), inset values show negative contributions of neonate (age 0) survival (*p*_*0*_) and infant survival (*p*_*1*_) that exceed axis limits.

**S7 Fig. Summed contributions for each population**. Stacked bars show summed contributions of infant, child and adult survival and of early, prime and late fertility. Positive and negative contributions are summed separately to reflect the net difference in population growth rate (white bars), relative to the composite mean hunter-gatherer reference. The black-and-white line crossing the bars indicates the population growth rate (*r* = log(*λ*)). Results are shown for ten small-scale societies and five chimpanzee populations (labeled as in Figure 2).

**S8 Fig. Total effects of vital rate differences**. Total effects (**S** *C*_*ij*\*_) reflecting the combined magnitude of contributions within each population (labeled as in Figure 2). Stacked bars decompose the total effect (the proportion of the combined magnitude of all contributions) made by infant, child and adult survival and by early, prime and late fertility (inset text shows the percent of total effects).

**S1 Table. Study populations and metadata**. Human populations are arranged by subsistence type (H: hunter-gatherers, A: acculturated hunter-gatherers, F: foraging-horticulturalists, P: pastoralists) and chimpanzees are arranged into wild (W), managed (M) and captive (C) populations. Columns contain information on location, region or continent, habitat type, population size, study period and data sources for fertility and mortality rates.

**S2 Table. Differences between populations**. For tests of differences in mean population growth rate (*r* = log *λ*), mortality measures, fertility measures, or selection measures (units above first row of values), rows show difference tests (*p, U*) for all humans vs. chimpanzees (*H*.*s*. vs. *P*.*t*.), for hunter-gatherers vs. chimpanzees (HG vs. *P*.*t*.), for non-foragers vs. chimpanzees (NF vs. *P*.*t*.), and for hunter-gatherers vs. non-foragers (HG vs. NF) Lower rows show means (x^-^) and standard deviations (SD) of each measure hunter-gatherers (HG), non-foragers (NF) or all humans (*H*.*s*.) and across declining (WC-) or increasing populations (WC+) of wild chimpanzees or across all wild chimpanzees (*P*.*t*.). NS results are indicated by greyed cells (*p* > 0.1) and marginal significance (*p* < 0.1) by lightly greyed cells; bold values indicate comparisons in which a given measure is significantly higher among humans (or hunter-gatherers) vs chimpanzees or among hunter-gatherers vs. non-foragers.

**S3 Table. Differences within populations**. Rows show Mann-Whitney-Cox *U*-statistics and *p*-values for differences in mean values between five hunter-gatherer societies (HG, n = 5), five non-exclusive forager societies (NF, n = 5), all ten human populations (*H*.*s*., n = 10), or five wild chimpanzee populations (*P*.*t*., n = 5). Columns to the left show results for tests of differences in either the total effect (combined magnitude of contributions) made by pairwise differences between total survival (*C*_*s*_), child survival (*C*_*c*_), adult survival (*C*_*a*_) and fertility (*C*_*f*_), or for differences in the relative accuracy of elasticities in predicting the fitness importance of child survival (*Z*_*c*_) vs. either adult survival (*Z*_*a*_) or fertility (*Z*_*f*_). Columns to the right report means (*x*^-^) and standard deviations (SD) for each of these composite measures across a given set of populations. NS results are indicated by greyed cells (*p* > 0.1) and marginal significance (*p* < 0.1) by lightly greyed cells; bold values indicate comparisons in which the first in a given pair of measures is significantly higher.

**S5 Table. Variable definitions**. Columns contain the variable symbol, variable name and source equation for the demographic parameters estimated in our analyses.

**S1 File. Methods details**. (1) Population metadata and ethnographic details; (2) Mortality smoothed with a Siler model; (3) Fertility smoothed with Loess regression; (4) Matrix model construction and emergent life history traits (5) Force of selection: vital rate sensitivities and elasticities; (6) Life table response experiments (LTRE): vital rate fitness contributions.

**S2 File. Results details**. (1) Population growth and decline; (2) Mortality and fertility patterns.

**S3 File. Supplemental References**. References supporting Supplemental information files.

